# Analysis of Copy Number Loss of the ErbB4 Receptor Tyrosine Kinase in Glioblastoma

**DOI:** 10.1101/154815

**Authors:** DeAnalisa C. Jones, Adriana Scanteianu, Matthew DiStefano, Mehdi Bouhaddou, Marc R. Birtwistle

## Abstract

Current treatments for glioblastoma multiforme (GBM)—an aggressive form of brain cancer—are minimally effective and yield a median survival of 14.6 months and a two-year survival rate of 30%. Given the severity of GBM and the limitations of its treatment, there is a need for the discovery of novel drug targets for GBM and more personalized treatment approaches based on the characteristics of an individual’s tumor. Most receptor tyrosine kinases—such as EGFR—act as oncogenes, but publicly available data from the Cancer Cell Line Encyclopedia (CCLE) indicates copy number loss in the ERBB4 RTK gene across dozens of GBM cell lines, suggesting a potential tumor suppressor role. This loss is mutually exclusive with loss of its cognate ligand NRG1 in CCLE as well, more strongly suggesting a functional role. The availability of higher resolution copy number data from clinical GBM patients in The Cancer Genome Atlas (TCGA) revealed that a region in Intron 1 of the ERBB4 gene was deleted in 69.1% of tumor samples harboring ERBB4 copy number loss; however, it was also found to be deleted in the matched normal tissue samples from these GBM patients (n = 81). Using the DECIPHER Genome Browser, we also discovered that this mutation occurs at approximately the same frequency in the general population as it does in the disease population. We conclude from these results that this loss in Intron 1 of the ERBB4 gene is neither a *de novo* driver mutation nor a predisposing factor to GBM, despite the indications from CCLE. A biological role of this significantly occurring genetic alteration is still unknown. While this is a negative result, the broader conclusion is that while copy number data from large cell line-based data repositories may yield compelling hypotheses, careful follow up with higher resolution copy number assays, patient data, and general population analyses are essential to codify initial hypotheses.

## INTRODUCTION

The ERBB/HER family of receptor tyrosine kinases (RTK’s) includes EGFR/ERBB1/HER1, ERBB2/HER2, ERBB3/HER3, and ERBB4/HER4^1-5^. Their activation by ligand binding followed by homo- and hetero-dimerization leads to activation of multiple mitogenic and survival pathways, such as the MAPK signaling pathway, which can drive cell proliferation and cell survival^1-3,5^. It is known that amplification in the gene copy number of ERBB/HER genes leads to overexpression and the sustained cell proliferation and survival in many cancers^1,2,4,5^. Overexpression of EGFR has been observed in many primary tumor types including lung, pancreas, breast, and glioblastoma while overexpression of HER2 has primarily been observed in breast and ovarian cancers^1,2,4^. Mutations such as these can be exploited when developing targeted cancer therapies^1-3,5^. For example, because EGFR is known to be overexpressed in non-small cell lung cancer (NSCLC), the most common type of lung cancer^5,6^, and NSCLC is treated using gefitinib—a kinase inhibitor that binds to the intracellular tyrosine kinase domain of EGFR and inhibits the signaling that drives cell proliferation and survival^6^. Amplification in HER2 copy number is observed in 20-30% of breast carcinomas^1^. Patients with this copy number variation are treated with trastuzumab—a monoclonal antibody that binds to the extracellular domain of HER2 to inhibit signaling that drives cell proliferation and survival^1,6^.

While much information is available about EGFR and HER2 signaling in cancer, less is known about the role of ERBB4. ERBB4 binds several ligands including betacellulin, HB-EGF, and epiregulin that also bind to EGFR^2^ but additionally other ligands such as neuregulin 1-4 (NRG1, NRG2, NRG3, and NRG4)^7^. ERBB4 is essential to cardiac, mammary, and neural development^2,8^ and is implicated in schizophrenia^9^. With regards to signaling and cancer, ERBB4 activates several of the same downstream proteins as EGFR—such as CBL, STAT5, and SHC^2^ but also strongly activates PI3K signaling^2,3^. Its overexpression has been associated with melanoma, medulloblastoma, and breast cancer progression^4,8^. However, it remains unclear what role, if any, ERBB4 plays in the progression of gliomas.

Copy number variations (CNV’s)—commonly termed deletions or amplifications—are generally accepted to be any genomic variations greater than 50 bp in length that alter the amount of DNA content of a gene^10,11^. CNV’s can play an important role in human disease—by altering the structure or abundance of transcripts and proteins, for example—or can have no phenotypic effect^10,11^. Examples in many human cancers include copy number loss of the gene that codes for the tumor suppressor protein PTEN^4,12^ and copy number gain in the gene that codes for the proto-oncogene EGFR^2^. Loss of a tumor suppressor gene, such as PTEN, and amplification of a protooncogene, such as EGFR, both lead to cancer progression but through different mechanisms ^1,2,4,12^. When a tumor suppressor is lost, it is no longer able to quell cell proliferation or induce cell death ^12^; and when a proto-oncogene is gained, amplified cell proliferation or inability to induce death occurs^1,2^. While a large volume of publicly available copy number data is generated using microarray and next-generation sequencing technologies^13,14,15^, much work remains in processing this data and interpreting the functional impacts of specific CNV’s in human disease^10^.

Here, we use publicly available data to explore copy number variation of ERBB4 in gliomas. Data from the Cancer Cell Line Encyclopedia (CCLE) suggests copy number loss of ERBB4 may be significant in glioma. However, subsequent follow up in The Cancer Genome Atlas (TCGA) and the DECIPHER Genome Browser demonstrate that the CCLE indications seem to be artifacts, which may be due to a combination of cell line models and low resolution copy number variation measurements. Regardless, this paper outlines a comprehensive approach to using publicly available copy number data to gain insight into the potential functional impact of CNV’s in cancer.

## METHODS

### Data Curation

*DNA Copy Number (41.6GB) Affy SNP* data in the form of copy number by gene for 60 glioma cell lines was downloaded from the Cancer Cell Line Encyclopedia (CCLE) data portal at https://portals.broadinstitute.org/. *Copy Number Segment* data from the *Affymetrix SNP 6.0* platform for 526 GBM tumor and matched normal tissue samples was downloaded from The Cancer Genome Atlas (TCGA) data portal at https://portal.gdc.cancer.gov. All downloaded data from CCLE and TCGA can be found in **Supplementary Data**. Copy number for the general (control) population was taken from the *Population: Copy-Number Variants* Affy6 consolidated data set of the DECIPHER Genome Browser and was accessed by chromosome location query. Copy number data from all three databases was generated using the Affymetrix SNP 6.0 microarray system.

### Data Processing

Raw microarray data describes each segment of the genome by a chromosome number and base pair range and a segment mean is assigned to each. This raw data can be converted to copy number by gene by locating the segment of the genome that contains the gene and then using the following formula to convert the segment mean (SM) to copy number (CN)^16^:

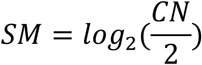

Downloaded data from CCLE was already converted into copy number by gene. Downloaded data from TCGA needed to be converted to copy number by gene, which was done using the above equation. All gene locations within the genome were taken from the UCSC Genome Browser on Human Feb. 2009 (GRCh37/hg19).

## RESULTS

### Gene Copy Number Distribution in CCLE

A long-term goal in our lab is to construct mechanistic mathematical models of glioma cell signaling that integrate commonly mutated signaling pathways and are tailored to an individual tumor’s genomic, transcriptomic, and proteomic data—such as the model proposed in Bouhaddou et al^17^. Public databases are a potentially valuable source of data for this research direction.

With this goal in mind, an initial analysis of copy number data from 60 glioma cell lines from the CCLE database revealed that the copy number distribution for ERBB4 is shifted to the left of normal copy number of 2 for diploid cells, signifying copy number loss or deletion (**Figure 1**). EGFR and PTEN were used as positive and negative controls respectively, and ERBB4 copy number is similar to the known tumor suppressor PTEN. This result is counter to what might be expected from a member of the ERBB family—as we can see from the right-shifted EGFR copy number distribution—and suggested that ERBB4 may be acting as a tumor suppressor in gliomas.

**Figure 1.**
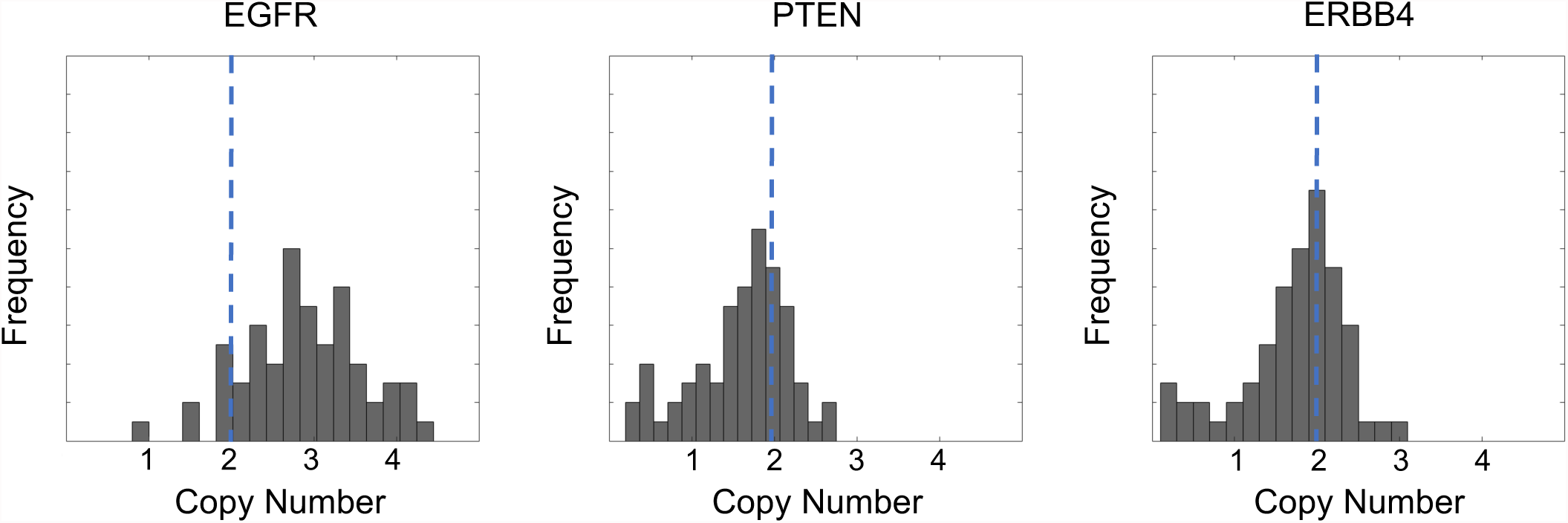
Gene Copy Number Distribution in CCLE. Gene copy number distributions of the ERBB4, EGFR, and PTEN genes across all glioma cell lines in CCLE (n = 60). The ERBB4 gene copy number distribution is shifted to the left of normal copy number of 2 for diploid cells, signifying copy number loss or deletion. Copy number data for EGFR and PTEN was analyzed to confirm what distribution properties are expected of a known oncogene (EGFR) and a known tumor supressor (PTEN). An oncogene, which is amplified in cancer, exhibits a right-shifted distribution while a tumor suppressor, which is lost in cancer, exhibits a left-shifted distribution.

### Mutually Exclusive Copy Number Loss of ERBB4 and NRG1 Genes

If ERBB4 is behaving as a tumor suppressor, only the receptor or the ligand—but not both—needs to be missing in order for there to be loss of tumor suppressor activity i.e. cancer progression. Therefore, we asked whether there was mutual exclusivity in copy number loss between ERBB4 and its endogenous ligand in the central nervous system, neuregulin-1 (NRG1). Copy number analysis of all glioma cell lines in CCLE revealed that there is a potential mutually exclusive relationship between loss in ERBB4 and loss in NRG1 as ERBB4 and NRG1 copy number loss never occur simultaneously (**Figure 2a-b**).

**Figure 2.**
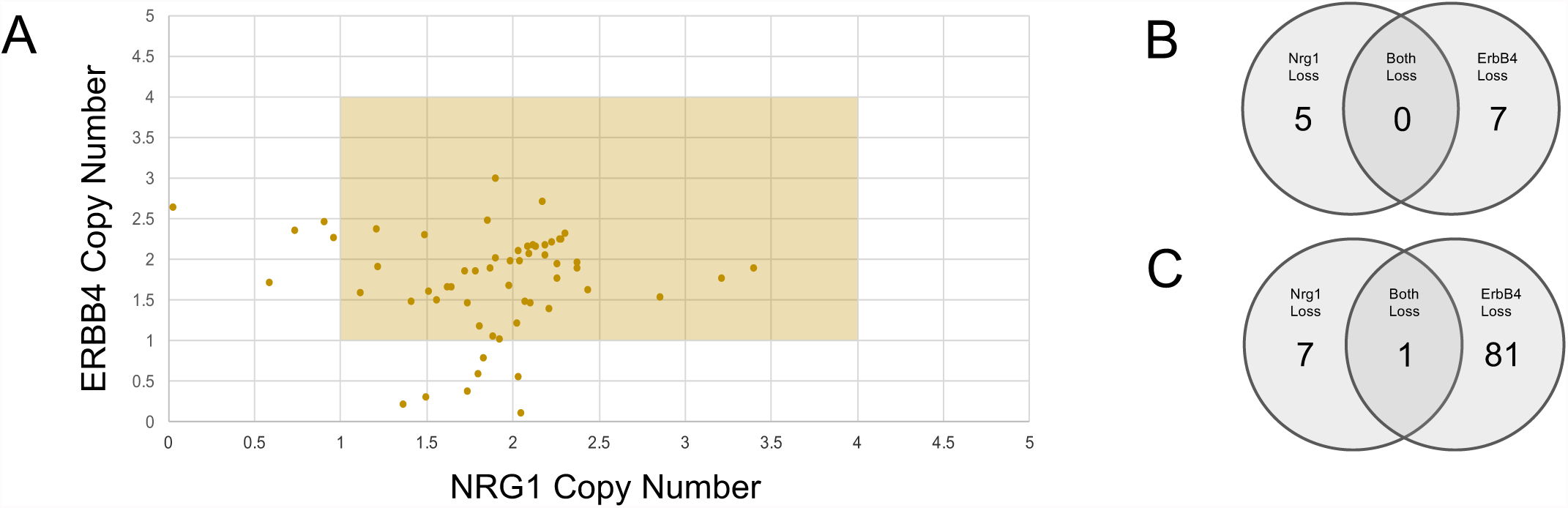
Mutually Exclusive Copy Number Loss of ERBB4 and NRG1 Genes. **(a)** Shown is ERBB4 copy number versus NRG1 copy number for each cell line where the highlighted region represents diploidy. Across all glioma cell lines in CCLE, ERBB4 and NRG1 copy number loss never occur simultaneously. **(b)** Across all glioma cell lines in CCLE (n = 60), ERBB4 copy number loss occurs in 7 cell lines and NRG1 occurs in 5 cell lines. Again, loss of both genes never occurs in the same cell line. **(c)** Across all GBM tumor samples from TCGA (n = 526), ERBB4 copy number loss occurs in 81 patients and NRG1 occurs in 7 patients. Both genes are only lost in 1 patient.

### ERBB4 Copy Number Loss in TCGA

Our preliminary analyses in glioma cell lines were expanded to GBM tumor samples from patients in TCGA to address whether or not the results from cell lines translate to clinical GBM patients. A mutually exclusive relationship was observed between ERBB4 and NRG1 copy number loss; however, loss of ERBB4 is strongly favored compared to NRG1 loss, as opposed to more parity in CCLE. While NRG1 loss was observed in only 7 of the tumor samples from 526 GBM patients in TCGA, ERBB4 loss was observed in 81 samples (**Figure 2c**). This result shifted our focus to copy number loss in the ERBB4 gene only in GBM tumor samples. When compared to EGFR and PTEN loss in GBM tumor samples, which again were used as positive and negative controls, the 15.4% frequency of ERBB4 copy number loss behaves similarly to that of GBM tumor suppressor PTEN (**Figure 3a**).

**Figure 3.**
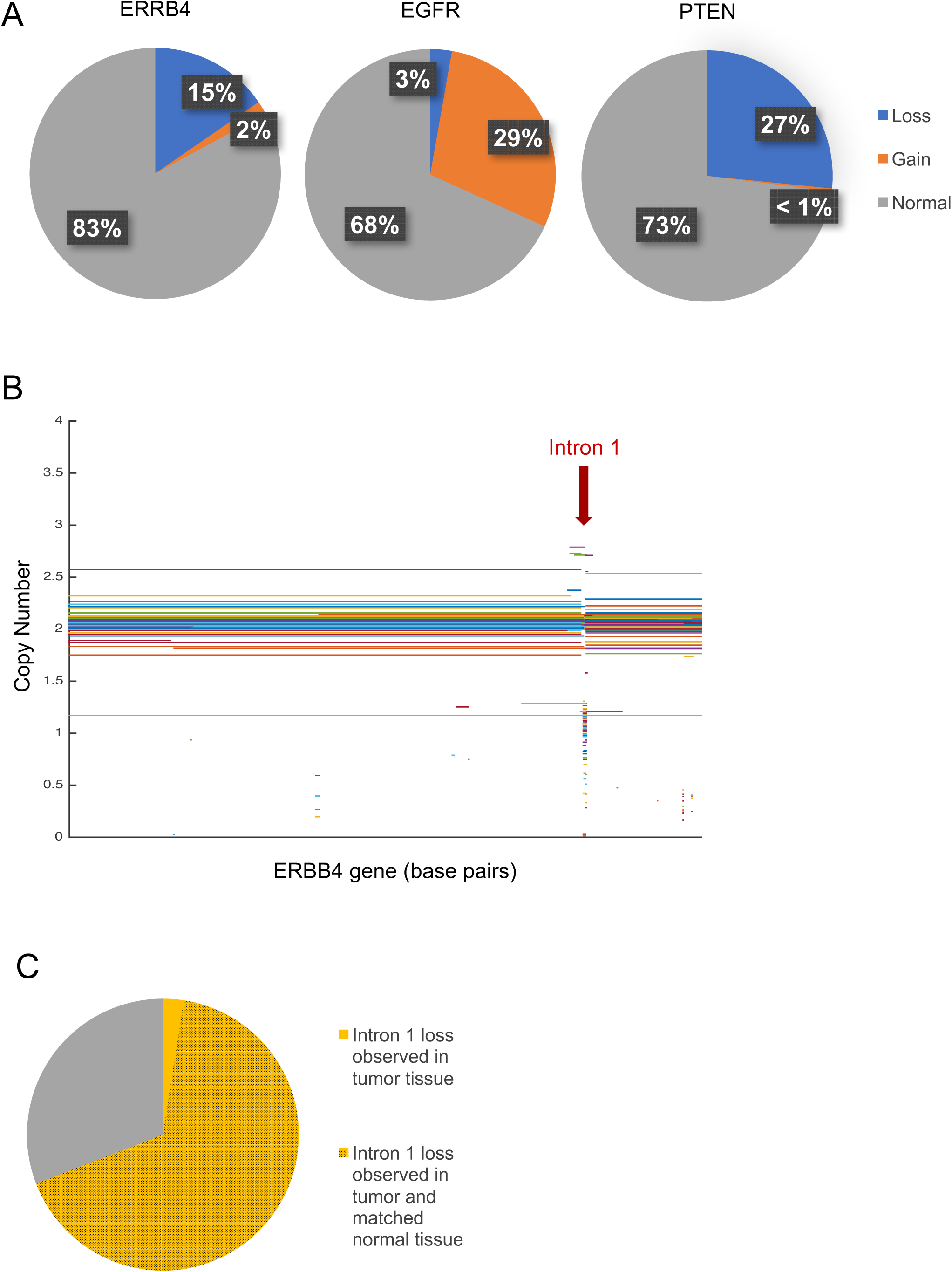
Frequency of Copy Number Loss in TCGA. **(a)** Shown are the percentages of normal copy number, copy number loss, and copy number gain observed for ERBB4, EGFR (a known oncogene in GBM), and PTEN (a known tumor suppressor in GBM) in GBM tumor samples from TCGA (n = 526). The ERBB4 gene was lost in 15.4% of samples. When compared to EGFR and PTEN, this frequency of loss behaves similarly to that of known GBM tumor suppressor PTEN. **(b)** Here, copy number values from segmented copy number data are mapped to segments of nucleotides within the ERBB4 gene. Copy number loss in ERBB4 appears to be localized to one region located in Intron 1 of the gene. **(c)** Of the tumor samples where ERBB4 loss was observed (n = 81), 69.1% of the loss was observed in a localized region within intron 1 of the ERBB4 gene. Of the tumor samples where loss in intron 1 occurred (n = 56), 96.4% of the matched normal tissue samples also demonstrated loss in intron 1.

Segmented copy number data available in TCGA allows us to localize copy number variations not only to whole genes but also to segments of nucleotides within genes. Comparing copy number to segments of nucleotides within the ERBB4 gene revealed that copy number loss in ERBB4 seems to be localized to one 5 kb region located in Intron 1 (**Figure 3b**). In fact, of the 81 GBM tumor samples where ERBB4 loss was observed, 69.1% of the loss was observed in this specific region. While this suggested that ERBB4 may demonstrate tumor suppressor activity that is compromised when a 5 Kb region in Intron 1 is deleted, we found that 96.4% of the matched normal tissue samples for these patients also demonstrated copy number loss in intron 1 (**Figure 3c**). Thus, this ERBB4 CNV is likely not a *de novo* somatic mutation that is a driver of GBM.

### Frequency of ERBB4 Copy Number Loss in the General Population

Although the clinical data do not support the initial suggestion from CCLE data that ERBB4 copy number loss is associated with GBM, it may be possible that loss in intron 1 of the ERBB4 gene is a factor that increases the risk that an individual will develop GBM. To investigate the possibility that this CNV may still be a predisposing factor to GBM, its frequency was characterized in the general, non-GBM population using consolidated database ^18^. population data from the DECIPHER. Querying “ERBB4” in DECIPHER’s genome browser returns common copy number variants observed within this gene in the general population in the *Population: Copy-Number Variants* track. Different studies used to obtain copy number information for the general population are merged into this database and separated by study. We used data from the *Affy6* study only (n = 5919), which was generated using the same Affymetrix SNP 6.0 microarray platform as was used in CCLE and TCGA as a part of the Sanger Institute’s Wellcome Trust Case Control Consortium (WTCCC) study^19^.

The frequency of CNV in intron 1 of the ERBB4 gene compared to instances of CNV in the EGFR and PTEN genes in the general population is depicted in **Figure 4**. It was found that the CNV occurs at a similar frequency in the general population (12.5%) as it does in the GBM population (15.4%). Comparison to the frequency of EGFR and PTEN CNV’s in the general, non-GBM population confirms that *de novo* driver mutations do not occur at the same frequency in the general population as they do in the disease population. *De novo* mutations demonstrate little CNV in the general population and increased CNV in the disease population. From this result, we concluded that loss in the ERBB4 gene is not a predisposing factor to GBM.

**Figure 4.**
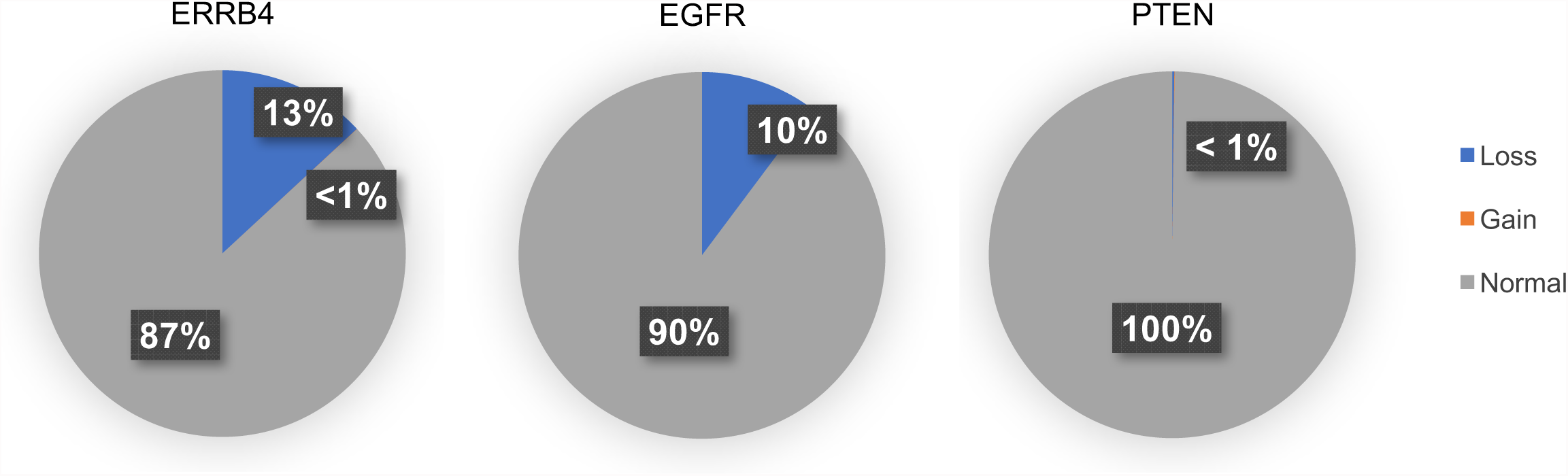
Frequency of Copy Number Loss in the General Population. Shown are the percentages of normal copy number, copy number loss, and copy number gain of the ERBB4 gene compared to known de novo driver mutations in GBM EGFR and PTEN in the general, healthy population from the DECIPHER database (n = 5919). Unlike in de novo driver mutations, the ERBB4 CNV we observed occurs at a similar frequency in the general population (12.5%) as it does in the GBM population (15.4%).

## CONCLUSIONS

A deeper investigation of copy number data in tumor tissue samples from GBM patients in TCGA and the general, healthy population in DECIPHER disproved our initial hypothesis that ERBB4 may be acting as a tumor suppressor in GBM. We attribute this artefactual initial finding from glioma cell lines in CCLE to a number of possible factors including the known limitations associated with both cell line studies^20^, the analysis of only gene-level and not segment-level data, as well as the lack of a matched normal control for cell lines. In addition, we noticed much variation in the resolution of base pair segments between patients and genes while analyzing copy number data generated using a microarray platform. For example, Patient 1 may have multiple ERBB4 copy number values because the ERBB4 gene spanned multiple segments in the microarray (i.e. higher resolution) while Patient 2 may have one copy number value that describes not only the ERBB4 gene but also other neighboring genes within the same chromosome because the microarray segment contained multiple genes (i.e. lower resolution). Use of whole exome sequencing technology to infer copy number may address copy number resolution issues mentioned here^21^. Nonetheless, our work offers a comprehensive methodology for using publicly available copy number data from CCLE, TCGA, and DECIPHER to infer the role of CNV’s in cancer progression.

## ACKNOWLEDGEMENTS

DeAnalisa Jones has received support from an NIGMS-funded diversity supplement to R01GM104184 (MRB) and the NIH Post-Baccalaureate Research Education Program (PREP) R25GM064118 grant.

## REFERENCES

1. Hudis, C. A. “Trastuzumab – Mechanism of Action and Use in Clinical Practice.” New England Journal of Medicine 357.1 (2007): 39–51.

2. Citri, A., and Y. Yarden. “EGF-ERBB Signalling: Towards the Systems Level.” Nature Reviews Molecular Cell Biology 7.7 (2006): 505–16.

3. Bowers, G., D. Reardon, T. Hewitt, P. Dent, R. B. Mikkelsen, K. Valerie, G. Lammering, C. Amir, and R. K. Schmidt-Ullrich. “The Relative Role of ErbB1-4 Receptor Tyrosine Kinases in Radiation Signal Transduction Responses of Human Carcinoma Cells.” Oncogene 20.11 (2001): 1388–397.

4. Davies, M. A., and Y. Samuels. “Analysis of the Genome to Personalize Therapy for Melanoma.” Oncogene 29.41 (2010): 5545–555.

5. Paez, J. G., P. A. Janne, J. C. Lee, S. Tracy, H. Greulich, S. Gabriel, P. Herman, F. J. Kaye, N. Lindeman, T. J. Boggon, K. Naoki, H. Sasaki, Y. Fujii, M. J. Eck, W. R. Sellers, B. E. Johnson, and M. Meyerson. “EGFR Mutations in Lung Cancer: Correlation with Clinical Response to Gefitinib Therapy.” Science 304.5676 (2004): 1497–500.

6. Gharwan, H., and H. Groninger. “Kinase Inhibitors and Monoclonal Antibodies in Oncology: Clinical Implications.” Nature Reviews Clinical Oncology 13.4 (2015): 209–27.

7. Ricci, C., L. Landuzzi, I. Rossi, C. De Giovanni, G. Nicoletti, A. Astolfi, S. Pupa, S. Menard, K. Scotlandi, P. Nanni, and P. L. Lollini. “Expression of HER/erbb Family of Receptor Tyrosine Kinases and Induction of Differentiation by Glial Growth Factor 2 in Human Rhabdomyosarcoma Cells.” International Journal of Cancer 87.1 (2000): 29–36.

8. Haskins, J. W., D. X. Nguyen, and D. F. Stern. “Neuregulin 1-activated ERBB4 Interacts with YAP to Induce Hippo Pathway Target Genes and Promote Cell Migration.” Science Signaling 7.355 (2014).

9. Bean, J. C., T. W. Lin, A. Sathyamurthy, F. Liu, D. M. Yin, W. C. Xiong, and L. Mei. “Genetic Labeling Reveals Novel Cellular Targets of Schizophrenia Susceptibility Gene: Distribution of GABA and Non-GABA ErbB4-Positive Cells in Adult Mouse Brain.” Journal of Neuroscience 34.40 (2014): 13549–3566.

10. Huang, N. The Functional Impact of Copy Number Variation in the Human Genome. Diss. Darwin College, U of Cambridge, 2011.

11. Zarrei, M., J. R. MacDonald, D. Merico, and S. W. Scherer. “A Copy Number Variation Map of the Human Genomes.” Nature Reviews Genetics 16 (2016): 172–83.

12. Chen, L., and D. Guo. “The Functions of Tumor Suppressor PTEN in Innate and Adaptive Immunity.” Cellular & Molecular Immunology (2017).

13. Barretina, J., G. Caponigro, N. Stransky, K. Venkatesan, A. Margolin, S. Kim, C. Wilson, J. Lehár, G. Kryukov, D. Sonkin, A. Reddy, M. Liu, L. Murray, M. Berger, J. Monahan, P. Morais, J. Meltzer, A. Korejwa, J. Jané-Valbuena, F. Mapa, J. Thibault, E. Bric-Furlong, P. Raman, A. Shipway, I. Engels, J. Cheng, G. Yu, J. Yu, P. Aspesi, M. de Silva, K. Jagtap, M. Jones, L. Wang, C. Hatton, E. Palescandolo, S. Gupta, S. Mahan, C. Sougnez, R. Onofrio, T. Liefeld, L. MacConaill, W. Winckler, M. Reich, N. Li, J. Mesirov, S. Gabriel, G. Getz, K. Ardlie, V. Chan, V. Myer, B. Weber, J. Porter, M. Warmuth, P. Finan, J. Harris, M. Meyerson, T. Golub, M. Morrissey, W. Sellers, R. Schlegel, and L. Garraway. “The Cancer Cell Line Encyclopedia Enables Predictive Modelling of Anticancer Drug Sensitivity.” Nature 483.7391 (2012): 603–07.

14. Bouhaddou, M., M. S. Distefano, E. A. Riesel, E. Carrasco, H. Y. Holzapfel, D. C. Jones, G. R. Smith, A. D. Stern, S. S. Somani, T. V. Thompson, and M. R. Birtwistle. “Drug Response Consistency in CCLE and CGP.” Nature 540.7631 (2016).

15. The Cancer Genome Atlas Research Network. “Comprehensive Genomic Characterization Defines Human Glioblastoma Genes and Core Pathways.” Nature 455 (2008): 1061–068.

16. “Copy Number Variation Analysis Pipeline.” Bioinformatics Pipeline: Copy Number Variation Analysis - GDC Docs. National Cancer Institute.

17. Bouhaddou, M., A. M. Barrette, R. J. Koch, M. S. Distefano, E. A. Riesel, A. D. Stern, L. C. Santos, A. Tan, A. Mertz, and M. R. Birtwistle. “An Integrated Mechanistic Model of Pan-Cancer Driver Pathways Predicts Stochastic Proliferation and Death.” BioRxiv (2017).

18. Firth, H. V., S. M. Richards, A. P. Bevan, S. Clayton, M. Corpas, D. Rajan, S. Van Vooren, Y. Moreau, R. M. Pettett, and N. P. Carter. “DECIPHER: Database of Chromosomal Imbalance and Phenotype in Humans Using Ensembl Resources.” The American Journal of Human Genetics (2009).

19. The Wellcome Trust Case Control Consortium. “Genome-wide Association Study of 14,000 Cases of Seven Common Diseases and 3,000 Shared Controls.” Nature 447 (2007): 661–78.

20. Balani, S., L. V. Nguyen, and C. J. Eaves. “Modeling the Process of Human Tumorigenesis.” Nature Communications 8.15422 (2017).

21. Landau, D. A., S. L. Carter, P. Stojanov, A. McKenna, K. Stevenson, M. S. Lawrence, C. Sougnez, C. Stewart, A. Sivachenko, L. Wang, Y. Wan, W. Zhang, S. A. Shukla, A. Vartanov, S. M. Fernandes, G. Saksena, K. Cibulskis, B. Tesar, S. Gabriel, N. Hacohen, M. Meyerson, E. S. Lander, D. Neuberg, J. R. Brown, G. Getz, and C. J. Wu. “Evolution and Impact of Subclonal Mutations in Chronic Lymphocytic Leukemia.” Cell 152.4 (2013): 714–26.

